# Intravital imaging of pulmonary lymphatics in inflammation and metastatic cancer

**DOI:** 10.1101/2024.09.12.612619

**Authors:** Simon J. Cleary, Longhui Qiu, Yurim Seo, Peter Baluk, Dan Liu, Nina K. Serwas, Jason G. Cyster, Donald M. McDonald, Matthew F. Krummel, Mark R. Looney

**Affiliations:** Department of Medicine, University of California, San Francisco (UCSF), CA, USA; Institute of Pharmaceutical Science, King’s College London, London, UK; Department of Anatomy, Cardiovascular Research Institute, and Helen Diller Family Comprehensive Cancer Center, UCSF, CA, USA; Howard Hughes Medical Institute and Department of Microbiology and Immunology, UCSF, CA, USA; Westlake Laboratory of Life Sciences and Biomedicine, Westlake University, Hangzhou, Zhejiang, China; Department of Pathology, UCSF, CA, USA; Bakar ImmunoX Initiative, UCSF, CA, USA

**Keywords:** Lymphatics, intravital microscopy, inflammation, dendritic cells, lung, pulmonary, leukocyte trafficking, cancer, metastasis

## Abstract

Intravital microscopy has enabled the study of immune dynamics in the pulmonary microvasculature, but many key events remain unseen because they occur in deeper lung regions. We therefore developed a technique for stabilized intravital imaging of bronchovascular cuffs and collecting lymphatics surrounding pulmonary veins in mice. Intravital imaging of pulmonary lymphatics revealed ventilation-dependence of steady-state lung lymph flow and ventilation-independent lymph flow during inflammation. We imaged the rapid exodus of migratory dendritic cells through lung lymphatics following inflammation and measured effects of pharmacologic and genetic interventions targeting chemokine signaling. Intravital imaging also captured lymphatic immune surveillance of lung-metastatic cancers and lymphatic metastasis of cancer cells. To our knowledge, this is the first imaging of lymph flow and leukocyte migration through intact pulmonary lymphatics. This approach will enable studies of protective and maladaptive processes unfolding within the lungs and in other previously inaccessible locations.

**Graphical abstract:** 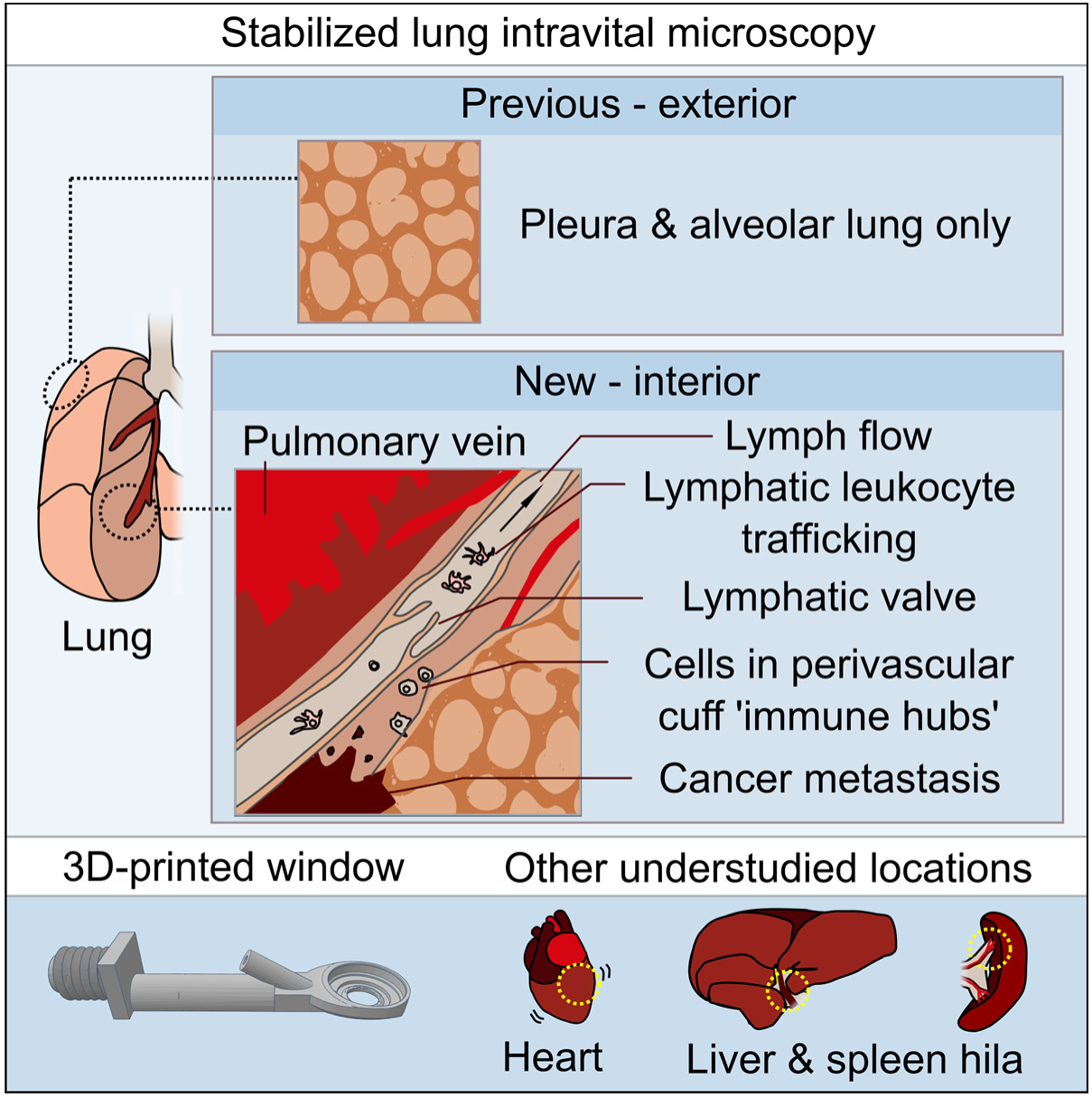

## Introduction

Stabilized intravital microscopy approaches have made it possible to directly study immune events that unfold within lung alveolar capillary units at subcellular resolution. Lung intravital microscopy has enabled mechanistic insights into lymphocyte surveillance (Looney et al., 2011; Podstawka et al., 2021), neutrophil recruitment (Looney et al., 2011; Conrad et al., 2022; Park et al., 2019), neutrophil extracellular trap release (Cleary et al., 2020; Lefrançais et al., 2018), platelet responses (Cleary et al., 2020, 2019), myeloid containment of lung-metastatic cancer cells (Headley et al., 2016), and alveolar macrophage patrolling (Neupane et al., 2020), as well as immune-modulatory and platelet-producing megakaryocytes in the lungs (Lefrançais et al., 2017; Pariser et al., 2021). Adapted lung imaging windows have permitted longer-term intravital imaging of events taking place over hours and days (Headley et al., 2016; Entenberg et al., 2018), and have also enabled imaging across outer surfaces of ventilated, perfused mouse lungs ex vivo (Banerji et al., 2023). However, all of these previous lung intravital microscopy approaches have been limited to alveolar lung tissue within ∼100 µm of distal pleural surfaces, a region devoid of important structures including major airways, large blood vessels and other structures of the lung interior.

The restriction of lung intravital microscopy to alveolar capillary units has prevented direct study of intact structures critical for pulmonary immune regulation. Notably, these understudied regions include bronchovascular cuff spaces that house unique leukocyte subsets and store reserves of edema fluid (Dahlgren and Molofsky, 2019). These spaces contain specialized lymphatics that transport fluid and cells out of the lungs and play vital but incompletely understood roles in lung fluid balance and immune responses in health and in various diseases (Trivedi and Outtz Reed, 2023). Intravital microscopy has proved useful for understanding function of other specialized blood and lymphatic vessels (Choe et al., 2015; Dixon et al., 2006; Collado-Diaz et al., 2022), but research into pulmonary lymphatic function has been limited by our inability to directly image intact lymphatics in the lungs (Trivedi and Outtz Reed, 2023; Baluk and McDonald, 2022; Stump et al., 2017).

We therefore developed novel tools and approaches that have enabled direct imaging of the movement of endogenous fluid and immune cells through intact lymphatics and cuff spaces surrounding pulmonary veins in the lungs of mice. We show that this approach can be used to answer key questions related to functions of lung lymphatic vessels in both draining fluid and in leukocyte trafficking during inflammatory responses and lung-metastatic cancer. In addition, apparatus and techniques developed for studying the lungs were also found to be useful for imaging other structures previously unseen using intravital microscopy. This article reports insights into pulmonary lymphatic biology using our new technique and provides a stabilization window model that can be 3D printed to allow other researchers to expand their studies to new tissue locations.

## Results

### An intravital microscopy approach enabling direct study of lung lymphatic function

The visceral pleural surfaces of lungs are accessible for intravital imaging but have few lymphatics in healthy mice, and those present in the exterior pleura are located far from the major sites of leukocyte and fluid trafficking in the lung interior (**Fig. 1A,B**) (Baluk et al., 2020; Yao et al., 2014). Direct observation of the dynamics of lymphatic valves, lymph flow and leukocyte trafficking in intact lymphatics in the lungs of living mice has therefore not been possible. Seeking an alternative location in the lungs to image lymphatics, we used cleared tissue imaging to image lymphatics across entire cleared lung lobes and observed that large collecting lymphatics follow pulmonary veins close to the proximal mediastinal surfaces of lungs (**Fig. 1A**). In addition to lymphatics, pulmonary veins are surrounded by cardiac muscle and perivascular cuff spaces that have both been implicated in immune regulation (Folmsbee et al., 2016; Dahlgren and Molofsky, 2019), so we developed an approach to stabilize and image these structures.

**Figure 1:**
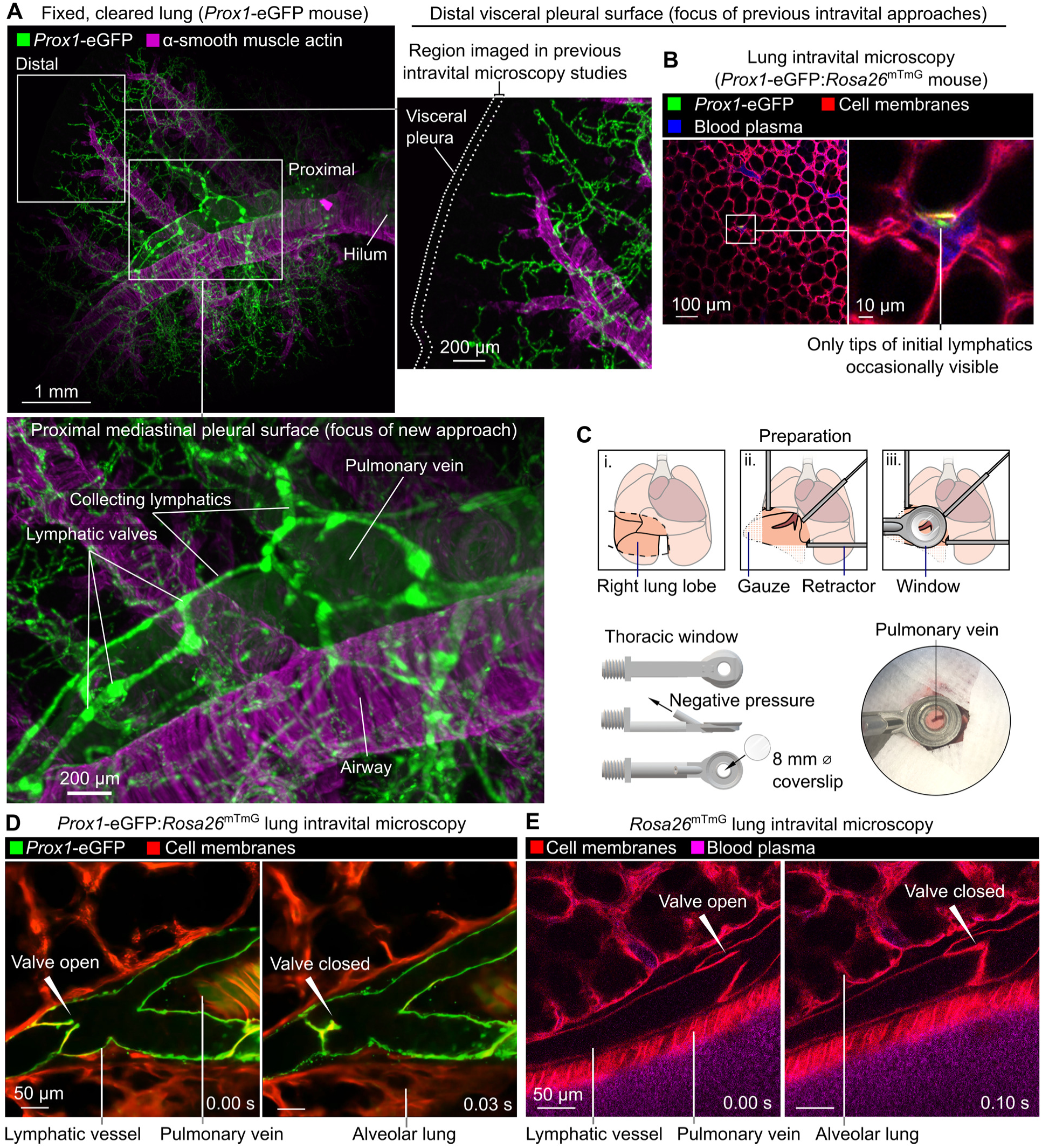
Intravital imaging of lymphatic vessels in lungs of ventilated, anesthetized mice. (**A**) Cleared lung from *Prox1*-eGFP mouse showing paucity of lymphatics near distal pleural surfaces and prominent collecting lymphatics surrounding pulmonary vein. (**B**) Distal lung intravital microscopy imaging of initial lymphatic tip in *Prox1*-eGFP:*Rosa26*^mTmG^ mouse. (**C**) Surgical preparation, window design and placement of window for imaging around pulmonary vein on mediastinal pleural surface. (**D**) Intravital imaging of functional lymphatic collectors in lungs of *Prox1*-eGFP:*Rosa26*^mTmG^ and (**E**) *Rosa26*^mTmG^ mice.

To immobilize areas around superficial pulmonary veins for intravital microscopy studies, we designed a 3D-printed stabilization window with a smaller frame than previous windows used for lung imaging (**Fig. 1C and Data File S1**) (Looney et al., 2011; Headley et al., 2016). This window was applied with a new surgical preparation to image previously unseen lung structures in ventilated, anesthetized mice expressing fluorescent reporters labelling lymphatic endothelial cells (*Prox1*-eGFP) (Choi et al., 2011) and all cell membranes (*Rosa26*^mTmG^) (Muzumdar et al., 2007) (**Fig. 1D**). We captured the opening and closing of pulmonary collecting lymphatic valves, pulmonary veins with pulsatile cardiac myocyte sheaths, and bronchovascular cuff spaces (**Fig. 1D and Video 1**). The distinctive bicuspid valves and bronchovascular cuff location of pulmonary collecting lymphatics enabled identification of these structures without a lymphatic-restricted reporter using *Rosa26*^mTmG^ mice (**Fig. 1E and Video 1**).

### Lymph flow and valve dynamics in intact lung lymphatics

Because pulmonary collecting lymphatics typically lack smooth muscle and pericyte coverage, they are thought to be unable to generate the intrinsic peristaltic contractions that drive lymph flow out from other organs (Outtz Reed et al., 2019). These anatomical features, together with evidence that changing respiratory rate has effects on thoracic duct outflow in large animal cannulation studies (Warren and Drinker, 1942), have led to the hypothesis that forces generated by ventilation primarily drive lung lymph flow. Intravital imaging of pulmonary collecting lymphatics allowed us to determine that, in steady state conditions with positive pressure ventilation, stabilized segments of pulmonary lymphatics do not display contractions but instead open and close their valves in synchrony with the respiratory rate (**Fig. 2A,B and Video 1**). Providing further evidence for a role for ventilation in driving steady state lung lymph flow, pausing mechanical ventilation resulted in cessation of pulmonary collecting valve opening and closing (**Fig. 2A-C and Video 1**). In contrast, one day after inducing acute lung inflammation by dosing bacterial lipopolysaccharides (LPS) into the lungs of mice, pulmonary lymphatic valves exhibited openings and closings that were asynchronous with ventilation and continued during ventilator pauses (**Fig. 2D-F and Video 1**). Tracking leukocytes that had entered lung lymph flow in LPS-treated mice, we confirmed that lymph flow out from inflamed lungs continues during ventilator pauses (**Fig. 2G,H and Video 1**). Together, these findings indicate that acute inflammation leads to uncoupling of lung lymph flow from ventilation, potentially driven by increased plasma extravasation from blood vessels made leaky by inflammation. These findings demonstrate the importance of studying pulmonary lymphatic biology in both normal physiology and in relevant disease models.

**Figure 2:**
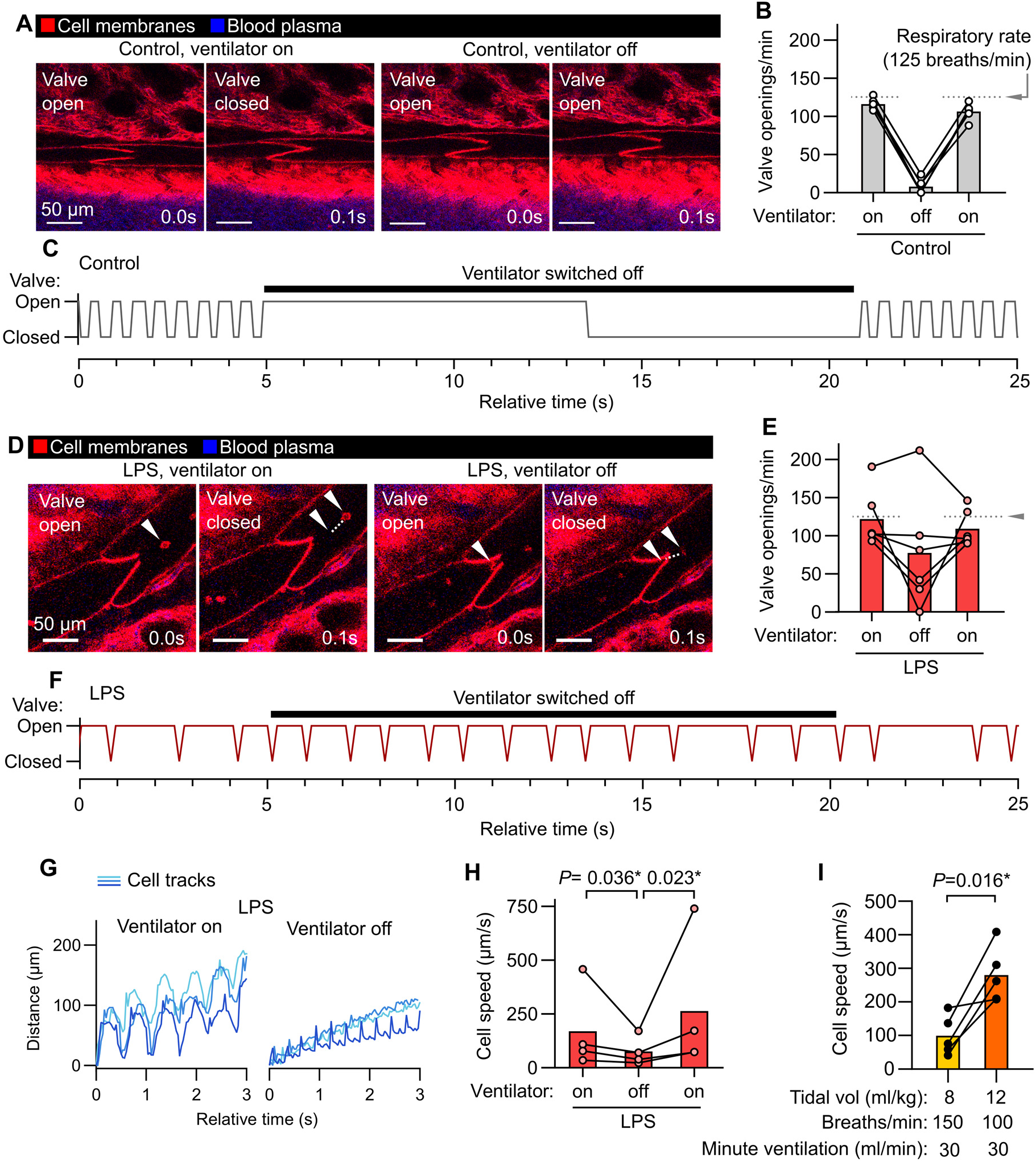
Ventilation-dependent and independent lymph flow through pulmonary collecting lymphatics. (**A**) Pulmonary lymphatic valves of steady-state control *Rosa26*^mTmG^ mice during ventilation and during a ventilator pause, with (**B**) quantification of effect on valve openings and (**C**) representative trace of valve status over time. (**D**) Pulmonary lymphatic valves from LPS-treated mice showing continuation of leukocyte flow and valve opening during ventilator pause, with (**E**) quantification of effect of ventilator pause on valve opening, (**F**) representative valve trace, and (**G**) representative traces of progress of tracked leukocytes through lymphatics with (**H**) quantification of speeds. (**I**) Cell speeds during lower versus higher tidal volume ventilation with indicated settings. Bar graphs show means, *P*-values are from: (**H**) repeated measures two-way ANOVA on log_10_-transformed data with Tukey’s multiple comparisons test; or (**I**) 2-tailed, paired t-test. Group sizes: (**B, E, I**) n=5; (**H**) n=4.

Mechanical ventilation with lower tidal volumes (6 ml/kg predicted body weight), compared to higher tidal volumes (12 ml/kg), decreases mortality in the acute respiratory distress syndrome (ARDS) (The ARDS Network, 2000). As lung inflammation changed the ventilation-dependence of lymph flow in inflamed lungs, and previous studies of the effects of tidal volume on lung lymph flow used ex vivo-perfused lungs from healthy sheep (Pearse et al., 2005), we examined the effect of ventilation with higher versus lower tidal volumes on lung lymph flow by directly imaging flow of native leukocytes in lymph leaving LPS-inflamed lungs. We compared ventilation with higher versus lower tidal volume using settings that matched minute ventilation. The higher tidal volume ventilation setting resulted in near-immediate increases in cell speeds in lymph flow (**Fig. 2I and Video 2**), highlighting coupling of pulmonary lymphatic function to lung distention and the utility of intravital microscopy for research into mechanisms of lung fluid balance.

### Leukocyte dynamics and diversity in lymphatics during lung inflammation

Previous intravital studies of lymphatics draining the skin and mesentery have revealed a stepwise process involving migration of leukocytes into lymphatic vessels (Pflicke and Sixt, 2009), followed by leukocyte crawling on the luminal lymphatic endothelial surface (Collado-Diaz et al., 2022), then leukocyte detachment for entry into lymph flow (Dixon et al., 2006). These events are important for adaptive immunity and immune tolerance, but have not been characterized using live imaging in intact lung lymphatics. Additionally, determining the cellular contents of lung lymph has been challenging using currently available approaches, particularly in small animals (Baluk et al., 2020; Ying et al., 1994; Tang et al., 2022; Stolley et al., 2020). Using our intravital imaging approach, we found that 24 hours after onset of LPS-induced lung inflammation, the majority of leukocytes in collecting lymphatics had entered lymph flow, moving at speeds of 25-500 µm/second (**Fig. 2I, 3A,B and Video 3**). Leukocytes were accompanied by lymphatic drainage of extravasated plasma protein, imaged using intravenously injected Evans blue dye (**Fig. 2A**). Live imaging lymphatics also revealed that lung lymphatics became distended in response to LPS (**Fig. S1**). Leukocytes were observed rolling on and becoming adhesive to the lymphatic endothelium (**Video 3**), indicating that, similar to the leukocyte adhesion cascade in blood vessels, a similar set of processes also enables immune surveillance within pulmonary lymphatics.

A large fraction of the cells entering lymph flow were dendritic cells with visible dendritic or veiled morphology, confirmed by imaging mice expressing the *Xcr1*-Venus reporter (labeling type 1 conventional dendritic cells) and *Itgax*-mCherry (labelling the majority of dendritic cells) (**Fig. 3C-E and Video 3**) (Cabeza-Cabrerizo et al., 2021). Monocyte/macrophage cells with high expression of Csf1r-eCFP reporter were also imaged within lymphatics (**Fig. 3E,F and Video 3**). Neutrophils have been observed in lymphatic vessels (Rigby et al., 2015; Lok et al., 2019), and using *MRP8*-Cre:*Rosa26*^mTmG^ neutrophil reporter mice we quantified neutrophil trafficking in pulmonary lymphatics after LPS treatment (**Fig. 3G,H and Video 3**). Using the *Pf4*-Cre:*Rosa26*^mTmG^ line, with labelling of megakaryocytes and platelets in the lungs, we observed only rare entry of platelet-sized particles into lung lymph flow following LPS treatment (**Fig. S2A**). As the *Itgax*-mCherry reporter also labels alveolar macrophages, and alveolar macrophages have been reported as trafficking to lung-draining lymph nodes but not imaged directly (Kirby et al., 2009), we labelled alveolar macrophages with PKH26 dye aggregates 5 days prior to imaging (Neupane et al., 2020), but did not observe alveolar macrophages entering lymphatics during the inflammatory response to LPS (**Fig. S2B**).

**Figure 3:**
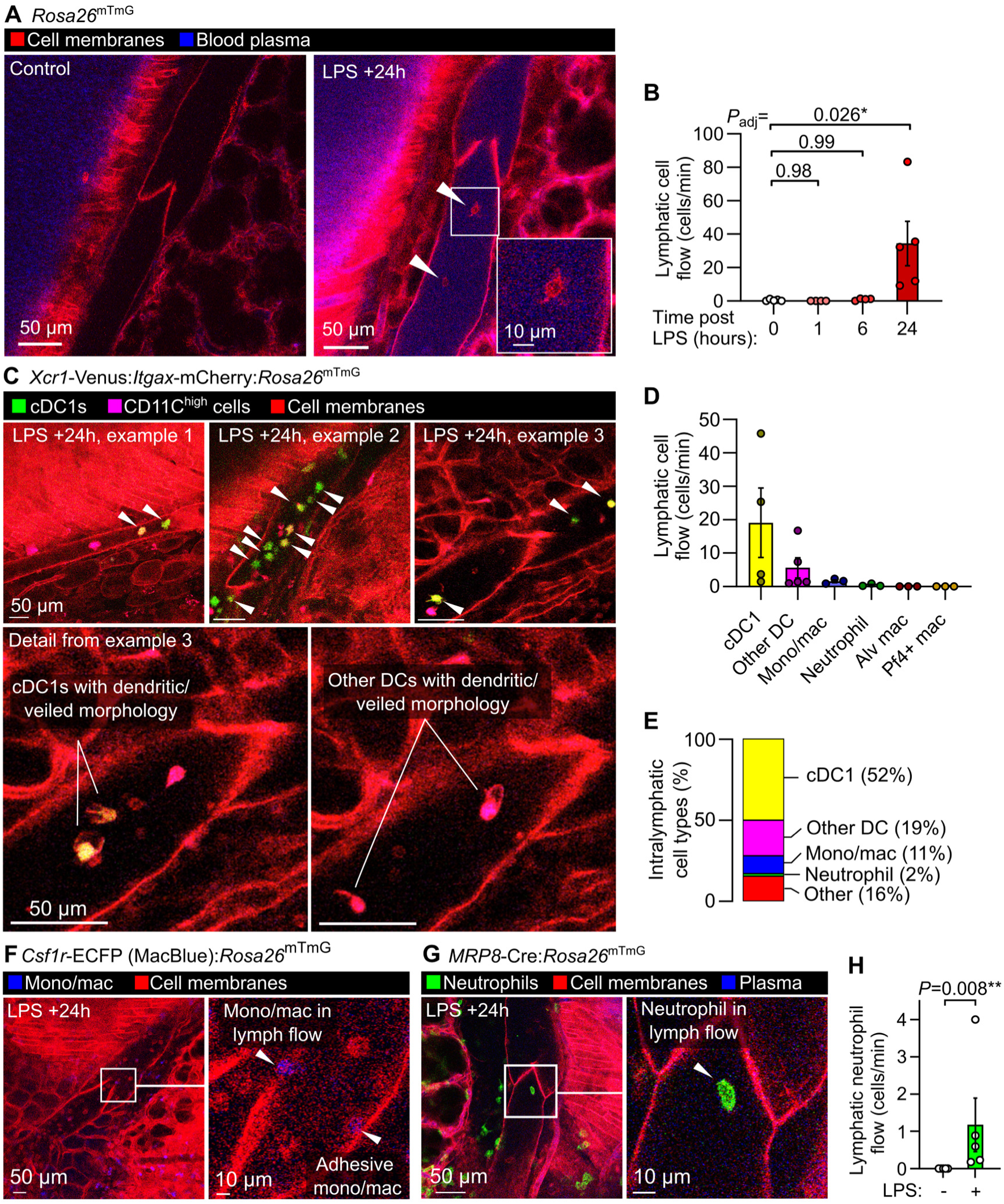
Dynamics and diversity of leukocyte trafficking within intact pulmonary lymphatics. (**A**) Pulmonary lymphatic vessels from a steady-state control and LPS-treated *Rosa26*^mTmG^ mice at 24 hours after onset of LPS-induced lung inflammation with arrowheads pointing to intralymphatic leukocytes. (**B**) Quantification of lymphatic flow of leukocytes. (**C**) Pulmonary lymphatics in *Xcr1*-Venus:*Itgax*-mCherry:*Rosa26*^mTmG^ mice at 24 hours after LPS treatment with arrowheads pointing to *Xcr1*-Venus+ cDC1s, with cell types in lymphatics quantified in (**D**) and (**E**) using the mouse lines shown in this figure and in Figure S2. Pulmonary lymphatic vessels from (**F**) *Csf1r*-ECFP:*Rosa26*^mTmG^ monocyte/macrophage reporter mouse or (**G**) *MRP8*-Cre:*Rosa26*^mTmG^ neutrophil reporter mouse 24 hours after LPS treatment with (**H**) quantification of lymphatic flow of neutrophils. Graphs show means ± SEM. *P*-values are from: (**B**) Kruskal-Wallis test with Dunn’s multiple comparisons to 0 hours (naïve) group; or (**H**) Mann-Whitney test. Group sizes: (**B**) n=4 (+1 and +6 hour groups), n=5 (0 and +24 hour groups); (**H**) n=5.

### Effects of interventions altering lymphatic trafficking of leukocytes

Mechanistically, G_αi_ protein-coupled receptors including Ccr7 and S1p1r have been implicated in lymphatic trafficking of leukocytes in mice (Hammad et al., 2003; Saeki et al., 1999; Czeloth et al., 2005). We inhibited signaling through G_αi_ subunits in mice using pertussis toxin, which eliminated lymphatic trafficking of immune cells in response to LPS inhalation (**Fig. 4A,B and Video 4**). *Ccr7* knockout leads to reduced leukocyte trafficking to lymph nodes, development of leukocyte aggregates in the lungs and defects in immune tolerance (Fleige et al., 2018). We confirmed that the bronchovascular cuff spaces in lungs *of Ccr7*^-/-^ mice become filled with leukocytes (**Fig. 4C**), and found that knockout of *Ccr7* greatly reduced leukocyte trafficking via pulmonary lymphatics one day after LPS treatment (**Fig. 4D,E and Video 5**).

**Figure 4:**
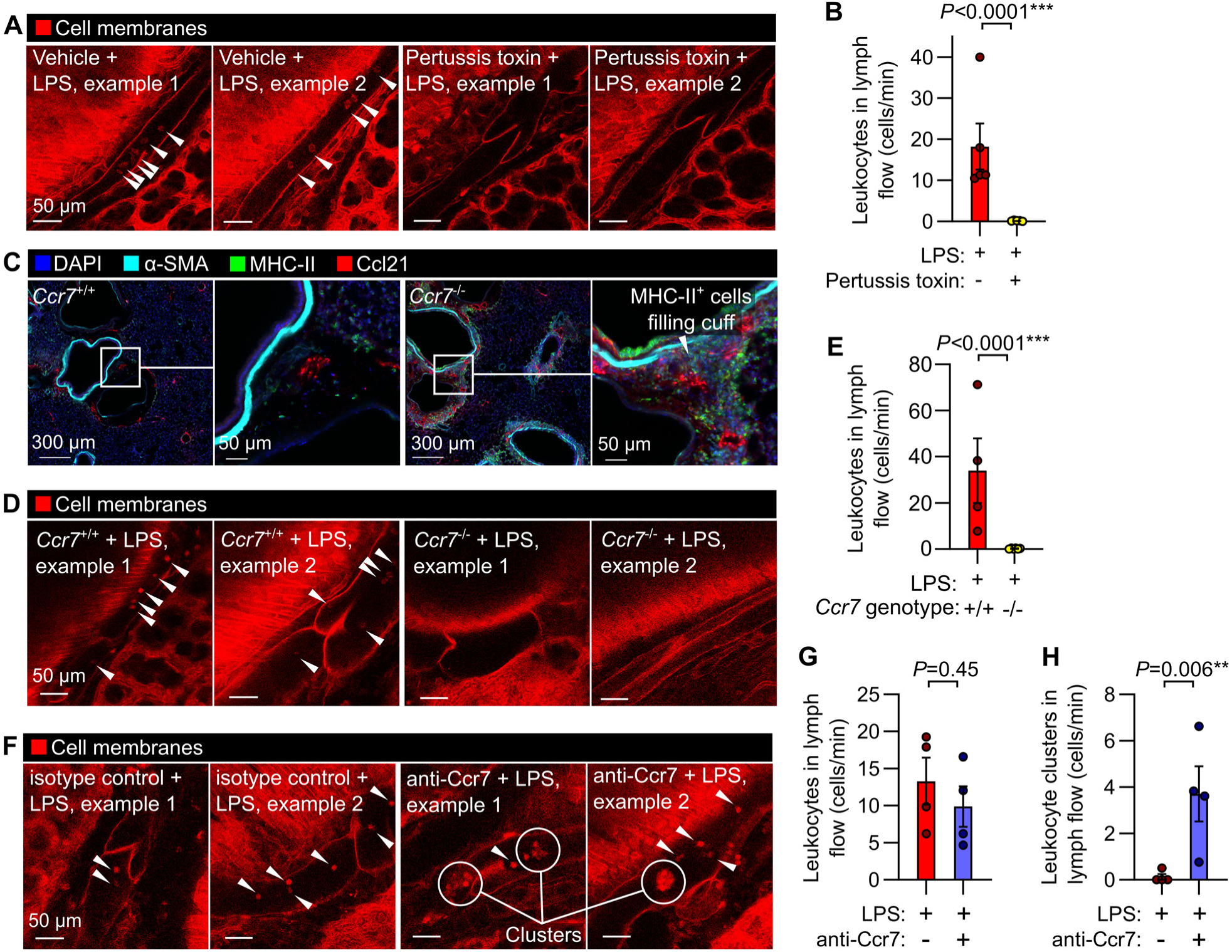
Interventions targeting chemokine receptor signaling alter leukocyte trafficking through lung lymphatics. (**A**) Leukocyte flow through pulmonary lymphatics from *Rosa26*^mTmG^ mice treated with either vehicle control or pertussis toxin before challenge with LPS with (**B**) quantification of effect of pertussis toxin. (**C**) Immunofluorescence images of *Ccr7*^+/+^ wild-type mice and *Ccr7*^-/-^ knockouts showing accumulation of MHC-II+ cells in bronchovascular cuffs. (**D**) Pulmonary lymphatic leukocyte flow in LPS-treated *Ccr7*^+/+^:*Rosa26*^mTmG^ mice and absence of intralymphatic leukocytes in *Ccr7*^-/-^:*Rosa26*^mTmG^ mice, with (**E**) quantification of effect of Ccr7 knockout. (**F**) Pulmonary lymphatics in LPS-treated *Rosa26*^mTmG^ mice given either isotype-matched control antibody or anti-Ccr7 by oropharyngeal aspiration together with LPS, with (**G**) absence of effect of antibody on total leukocyte flow and (**H**) quantification of flow of clusters of leukocytes in lung lymph. White arrowheads indicate intralymphatic leukocytes, circles highlight intralymphatic leukocyte clusters. Graphs show means ± SEM. *P*-values are from unpaired, two tailed t-tests on log_10_-transformed datasets. Group sizes: (**B**) n=5; (**E**, **G**, **H**) n=4.

Antibodies targeting human CCR7 are under clinical investigation, and antibodies targeting mouse Ccr7 have been used as research tools (Cuesta-Mateos et al., 2021; Liu et al., 2023; Pei et al., 2019). To understand how these agents might be altering pulmonary lymphatic function, we tested the effect of administering a functional Ccr7 blocking antibody, previously used for in vivo neutralization studies (Liu et al., 2023; Pei et al., 2019), on lymphatic leukocyte trafficking. After delivery of this Ccr7 blocking antibody directly into the lungs together with LPS, we found that Ccr7 blockade did not prevent entry of leukocytes into pulmonary collecting lymphatics but instead caused the appearance of large clusters of leukocytes that still achieved entry into lymph flow (**Fig. 4F-H and Video 6**). This discrepancy between the effects of constitutive genetic disruption of Ccr7 and blocking antibody treatment indicates that our understanding of constitutive versus induced loss of Ccr7 function is incomplete. These results highlight the usefulness of intravital lymphatic imaging for mechanistic studies of leukocyte trafficking through pulmonary lymphatics in inflammation.

### Lymphatic immune surveillance of metastatic tumors

Lymphatic vessels are also of great interest in cancer research because lymphatic-dependent immune responses, lymphatic metastasis, and lymphangiogenesis have been linked to altered cancer outcomes (Ma et al., 2018; Shields et al., 2010; Ubellacker et al., 2020; Steele et al., 2023). We therefore developed a protocol for imaging invasion of cancer cells and resultant immune surveillance responses in the lungs. We modeled lung metastasis by i.v. injecting *Rosa26*^mTmG^ mice with B16.F10 mouse melanoma cells engineered to express ZsGreen, a bright fluorophore that allows simultaneous imaging of entire tumor cells and their subcellular fragments. ZsGreen fluorescence also enables detection of cancer cell material taken up by immune cells because it retains fluorescence following phagocytosis (Ruhland et al., 2020). At 18 days after lungs were seeded with melanoma cells when pulmonary metastases are prevalent (**Fig. 5A)** (Ya et al., 2015), we observed leukocyte trafficking within lymphatics (**Fig. 5B,C and Video 7**). The majority (approximately two-thirds) of intralymphatic leukocytes contained material from cancer cells (**Fig. 5D**), indicating active lymphatic immune surveillance despite the failure of antitumor immune responses to clear cancer cells in this model without immunotherapy interventions (Ya et al., 2015). Intravital imaging also captured lymphatic metastasis of cancer cells (**Fig. 5B, E and Video 7**), revealed enrichment of bronchovascular cuff spaces with cancer cell material (**Fig. 5F,G**), and the frank invasion of collecting lymphatics by tumors (**Fig. 3H**). These results demonstrate the potential of this method for direct measurements of metastasis and tumor-immune interactions in lymphatics and interstitial spaces.

**Figure 5:**
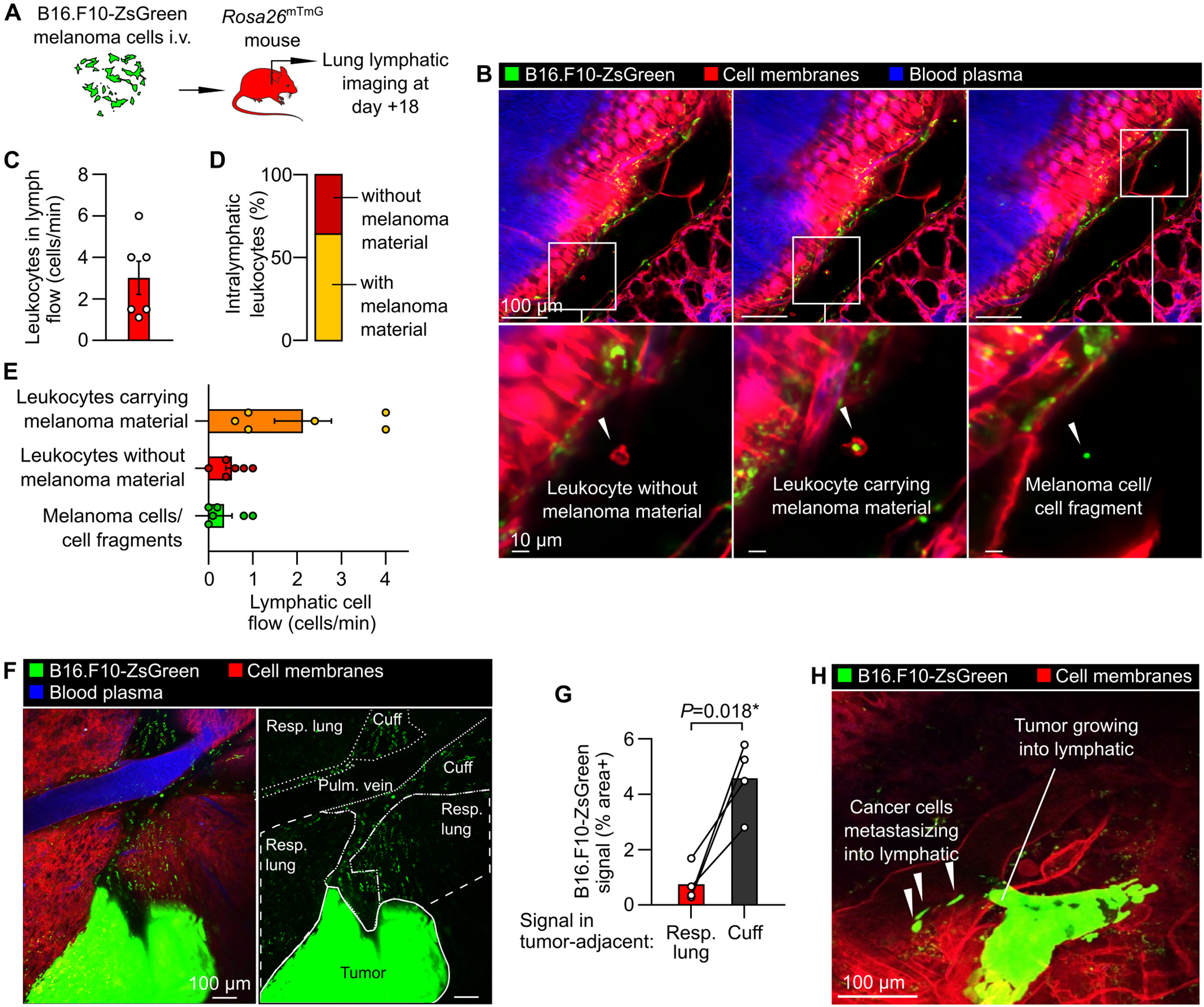
Tumor-immune interactions and metastasis within pulmonary lymphatics. (**A**) Schematic diagram summarizing B16.F10-ZsGreen melanoma mouse model. (**B**) Images of leukocytes carrying melanoma material, leukocytes without melanoma material and melanoma cell/cell fragments in lymph flow, with (**C**) quantification of total leukocyte flow and (**D, E**) breakdown of cell types observed in lymphatics. (**F**) Overview showing metastatic tumor in lung with (**G**) enrichment of bronchovascular cuff space (‘cuff’) relative to respiratory (‘resp.’, i.e. alveolar) lung. (**H**) Image showing intralymphatic tumor with melanoma cells detaching to enter lymph flow. Graphs show means ± SEM. *P*-value in (**G**) is from a 2-tailed, paired t-test. Group sizes: (**B**, **D**) n=6, (**F**) n=4.

### Imaging lymphatics draining other organs

Lastly, we tested whether our stabilization window could also be useful for imaging other tissues that are challenging to access and stabilize. With a similar approach, we imaged lymphatics in the hepatic hilum near the point of entry of the portal vein **(Fig. S3A)**. We imaged lymphatics draining the spleen, where leukocytes with lymphocyte morphology were abundant under normal conditions (**Fig. S3B**). In addition, the window developed in this study also enabled imaging of lymphatics within the beating heart (**Fig. S3C**). The stabilization approach reported in this study can therefore be used for intravital microscopy experiments in a diverse range of other understudied tissues and organs.

## Discussion

To our knowledge, the method reported in this study has enabled the first direct visualization of cellular dynamics within intact pulmonary lymphatics and bronchovascular cuff spaces. This new intravital microscopy approach solves several problems that have limited previous studies of pulmonary lymphatic function. Lung intravital microscopy has previously only been applied to the distal alveolar microvasculature, whereas this new method enables imaging of collecting lymphatics, bronchovascular cuff spaces and pulmonary veins, each of which has specialized and disease-relevant features that warrant direct study (Dahlgren and Molofsky, 2019; Trivedi and Outtz Reed, 2023; Baluk and McDonald, 2018, 2022). Approaches that have been established for studying pulmonary lymphatic function have involved excision of lungs, lymphatic vessels or lymph nodes, the cannulation of extrapulmonary lymphatics or microinjections into lung interstitial spaces (Outtz Reed et al., 2019; Dahlgren and Molofsky, 2019; Folmsbee and Gottardi, 2017). In our method, but not in these previous approaches, lymphatic function can be studied with continual ventilation, perfusion and innervation as well as intact flow through lymph nodes and thoracic duct outflow into the bloodstream. Use of genetically encoded fluorophores for monitoring cell trafficking and lymph flow also avoids potential artifacts from effects of tracer injections into delicate lung air or interstitial spaces, or ex vivo manipulation and adoptive transfer of cells.

The requirement of positive pressure ventilation is a limitation of our approach, although this feature is of relevance to the millions of people worldwide annually who receive supportive care from mechanical ventilation with conditions such as ARDS (Wunsch et al., 2010). Our stabilization approach requires application of gentle suction, but involves use of pressures that do not cause inflammation (Conrad et al., 2022; Cleary et al., 2020). We used mice in this study to facilitate use of gene modifications and interventions, but similar approaches will likely be useful in other model organisms, particularly as transgenic reporters are increasingly available in other animals, e.g. the *Prox1*-eGFP rat line (Jung et al., 2017). Our current preparations for lung imaging are limited to studying lymphatics running parallel to pulmonary veins. Whole-lung imaging (e.g. **Fig. 1A**) confirms that vein-associated collecting lymphatics receive lymphatic outflow from throughout the lungs, but it remains unclear whether pulmonary vein-associated collecting lymphatics differ from other collectors in the lungs.

Intravital microscopy studies using this method will be useful for investigating emerging concepts in lymphatic biology, including intralymphatic coagulation (Summers et al., 2022; MacDonald et al., 2022), lymphatic junctional plasticity (Baluk and McDonald, 2022; Churchill et al., 2022), induction of pulmonary lymphangiogenesis (Baluk et al., 2020; Szőke et al., 2021), as well as the incompletely understood role of lymphatics in major lung diseases including COVID-19, asthma, pulmonary fibrosis and tuberculosis (Trivedi and Outtz Reed, 2023). Beyond the lymphatic system, the versatile stabilization window that we developed for this study will also be useful for revealing unseen biology in a range of other tissues that are either difficult to access or challenging to image due to intrinsic motility.

## Methods

### Thoracic window production and assembly

Thoracic windows were 3D-printed in high-detail stainless steel using powder bed fusion (i.materialise, model provided as **Data File S1**). After polishing of the steel frame, an 8 mm #1 round coverslip (Thomas Scientific Cat# 64-0701) was inserted into the immersion liquid holder and sealed by using a needle to apply epoxy resin onto the outer edges of the coverslip and supporting steel surface. Following overnight drying, sealing of the coverslip onto the steel frame was confirmed by checking for retention of water added to the immersion liquid holder during aspiration through the suction port. Thoracic windows were cleaned with Terg-a-zyme (Sigma Aldrich Cat# Z273287), spraying with 70% ethanol and rinsing with sterile deionized water.

### Animal studies

Animal studies were conducted with approval from the UCSF institutional animal care and use committee. Male and female mice were used at age 6-16 weeks, and all mice were bred and maintained in the specific pathogen-free facility at UCSF. Prox1-eGFP mice (Choi et al., 2011) were from Donald M. McDonald (UCSF). *Rosa26*^mTmG^ mice were from Jax (Cat# 007576).(Muzumdar et al., 2007) Xcr1-Venus mice (Yamazaki et al., 2013), CD11c-mCherry mice(Khanna et al., 2010), and MacBlue mice (Sauter et al., 2014) were from Matthew F. Krummel (UCSF). MRP8-Cre(Passegué et al., 2004) and *Pf4*-Cre (Tiedt et al., 2007) mice were from Jax (Cat# 021614 and Cat# 008535, respectively). As previously, Evans blue dye (3 mg/kg, 0.75 mg/ml in 100 µl PBS) was injected i.v. immediately prior to imaging to label blood plasma proteins (Cleary et al., 2020), and PKH26-phagocytic cell linker was given by oropharyngeal aspiration (o.a.) as a 0.5 µM solution at 75 µl per mouse 5 days before imaging to label alveolar macrophages (Neupane et al., 2020). To induce acute lung inflammation we administered mice a single dose of LPS (O55:B5, Sigma-Aldrich Cat# L2880) at 4 mg/kg in PBS by o.a. dosing (Seo et al., 2023; Conrad et al., 2022). Pertussis toxin (Sigma-Aldrich Cat# P2980-50UG) was given i..v. immediately after LPS dosing at 1 µg per mouse in 100 µl PBS. Functional grade anti-Ccr7 clone 4B12 was purchased from Invitrogen (Cat# 50-144-95), compared to treatment with a non-reactive isotype-matched control clone 2A3 (BioXCell Cat# BE0089), given o.a. together with LPS at 50 µg per mouse in a total volume of 70 µl PBS.

### Intravital microscopy preparation for imaging the mediastinal visceral lung pleura

We anesthetized mice with ketamine/xylazine (60/40 mg/kg, i.p.), shaved their right chests and performed tracheal intubation for mechanical ventilation with room air containing 1% isoflurane at 10 µl/g body weight delivered at 125 breaths per minute with 2.5 cmH_2_O positive end expiratory pressure using a MiniVent system (Harvard Apparatus). Mice were then placed in the supine position, and an opening in the skin of the chest and underlying fascia was made to expose the right anterior ribcage. Ribs 2-4 were transected immediately to the right of the sternum and at posterior lateral locations and removed to make an opening in the ribcage, with point retractors placed to expose right lung lobes (**Fig. 1C**). The inferior right lobe was repositioned with a saline-moistened cotton-tipped applicator so that its mediastinal pleural surface faced upwards. The imaging window was then lowered over a pulmonary vein and application of negative pressure (-20 mmHg) was used to immobilize a segment of lung against the coverslip.

### Microscopy

For intravital microscopy we used a Nikon A1r microscope with a CFI75 Apochromat 25XC water immersion objective and high-frequency HD25 resonance scanner (UCSF Biological Imaging Development CoLab). Fluorescent excitation was achieved using a Mai Tai DeepSea IR laser (950 nm) for multiphoton imaging and, where required, Coherent OBIS lasers (405, 488, 561 and 647 nm), with emitted light filtered through 440/80, 525/50, 600/50 and 685/70 nm emission filters.

### Immunofluorescence

For 3D imaging of fixed lungs (4% formaldehyde by tracheal inflation and immersion overnight) we used CUBIC clearing with immunostaining for GFP (AlexaFluor 647-conjugated rabbit polyclonal, Invitrogen Cat# A-31852) and α-smooth muscle actin (αSMA, Cy3-conjugated clone 1A4, Sigma-Aldrich Cat# C6198) for imaging entire adult lung lobes, as previously described(Takahashi et al., 2022), with imaging on a customized light sheet microscope based around a Nikon AZ100 system with an AZ-Plan Apo 2x NA 0.2 objective and Vortran Laser Launch providing excitation at 561 and 640 nm and 605/52 and 705/72 emission filters (UCSF Center for Advanced Light Microscopy). For imaging lung sections 200 µm cryosections were prepared, stained and imaged as described in our previous work(Cleary et al., 2020, 2024). Primary antibodies used were: FITC-conjugated mouse anti-αSMA (clone 1A4, Sigma-Aldrich Cat# F37777); rat anti-MHC-II (clone M5/114.15.2, Invitrogen Cat# 16-5321-81) and goat anti-Ccl21 (R&D Systems Cat# AF457) with the latter two unconjugated antibodies detected using cross-adsorbed Donkey polyclonal secondaries: AlexaFluor 647-conjugated anti-rat IgG and Cy3-conjugated anti-goat IgG (Jackson Immunoresearch Cat# 712-605-153 and Cat# 705-165-147, respectively). Sections were imaged using the Nikon A1r confocal microscope described above.

### Metastatic melanoma model

As in previous reports(Headley et al., 2016; Ruhland et al., 2020; Ya et al., 2015), we gave *Rosa26*^mTmG^ mice an intravenous injection containing 1×10^5^ B16.F10-ZsGreen cells for seeding pulmonary melanoma metastases.

### Preparations for imaging other organs

Mice were anesthetized as described above. For spleen and liver imaging, organs were flipped in a cranial direction to expose hilar structures, with the window placed on the border of organs and interstitial tissue. For imaging the heart, similar to previous approaches but without use of glue(Lee et al., 2012), heart tissue was exposed with a left-side thoracotomy and placing of the stabilization window over the left ventricle.

## Acknowledgements

Microscopy work was possible due to support from the UCSF imaging facilities: the Biological Imaging Development CoLab (BIDC, with special thanks to Kyle Marchuk and Austin Edwards) and Center for Advanced Light Microscopy (CALM, with special thanks to SoYeon Kim).

## Author contributions

Conceptualization: S.J.C., L.Q., M.R.L.

Methodology: S.J.C., L.Q., Y.S., P.B., D.L., N.S., J.G.C., D.M.M., M.F.K., M.R.L.

Investigation: S.J.C., L.Q., M.R.L.

Funding acquisition: D.M.M., M.R.L.

Writing – original draft: S.J.C., M.R.L.

Writing – review & editing: S.J.C., L.Q., Y.S., P.B., D.L., N.S., J.G.C., D.M.M., M.F.K., M.R.L.

## Supplemental material

**Video 1** Effects of changing ventilator settings on pulmonary lymphatic valve function and afferent lung lymph flow

**Video 2** Effect of changing ventilator tidal volume on lymph flow within pulmonary lymphatics following LPS-induced acute lung inflammation

**Video 3** Leukocyte dynamics and diversity within pulmonary lymphatics after LPS-induced acute lung inflammation

**Video 4** Effect of pertussis toxin on leukocyte flow within pulmonary lymphatics following LPS-induced acute lung inflammation

**Video 5** Effect of knockout of *Ccr7* on leukocyte flow within pulmonary lymphatics following LPS-induced acute lung inflammation

**Video 6** Effect of Ccr7 blocking antibody treatment on leukocyte flow within pulmonary lymphatics following LPS-induced acute lung inflammation

**Video 7** Pulmonary lymphatic trafficking of leukocytes, cancer cell material and cancer cells following lung metastasis of B16.F10 melanoma cells

**Supplementary Data File 1** 3D model of thoracic window

---

**Figure S1:**
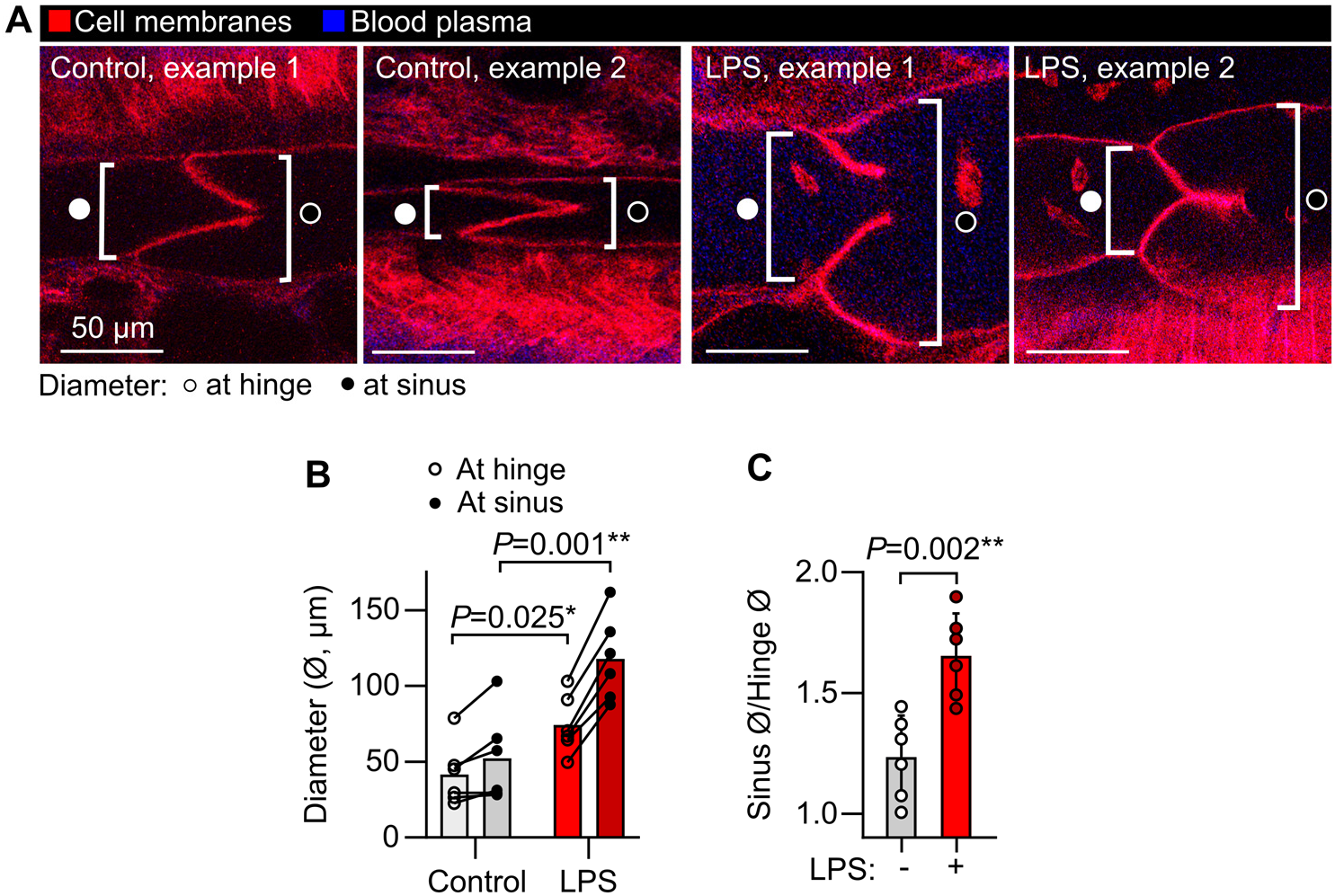
Measurement of pulmonary lymphatic distension in LPS-induced acute lung inflammation. (**A**) Representative images of pulmonary lymphatic valves from steady state controls and LPS-treated *Rosa26*^mTmG^ mice showing approach for measuring lymphatic diameter. (**B**) Lymphatic vessel diameters at valve hinges and at sinuses immediately downstream of valves. (**C**) Sinus diameters divided by hinge diameters showing relative distension of sinuses. Graphs show means ± SEM. *P*-values are from: (**B**) repeated measures, 2-way ANOVA with Holm-Šídák test for effect of LPS within vessel region groups; or (**C**) unpaired, 2-tailed t-test. Group sizes: n=6.

**Figure S2:**
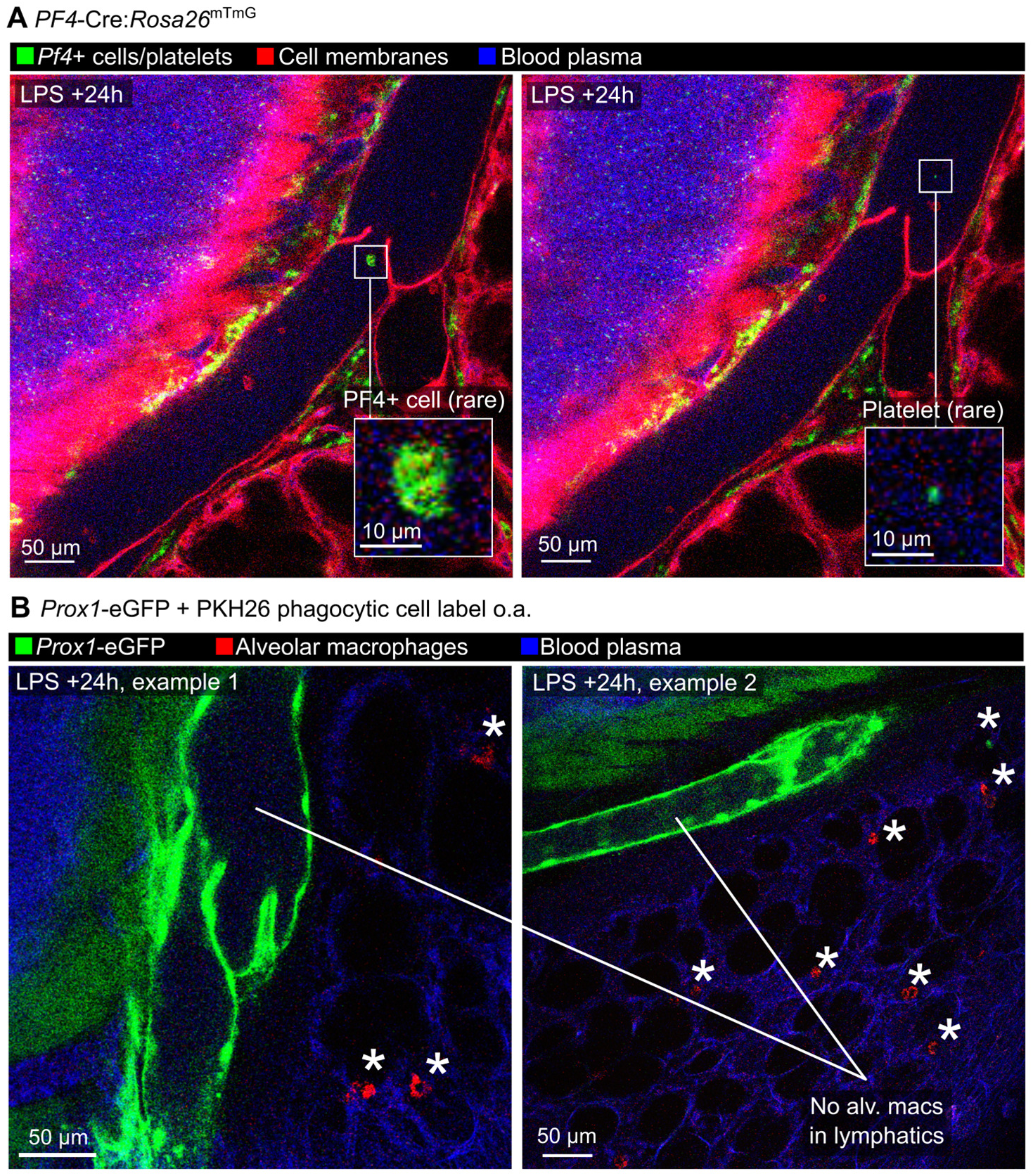
Imaging of pulmonary lymphatics in LPS-treated *Pf4*-Cre:*Rosa26*^mTmG^ mice and *Prox1*-eGFP mice given PKH26-PCL to label alveolar macrophages. (**A**) Intravital images of an LPS-treated *Pf4*-Cre:*Rosa26*^mTmG^ mouse showing platelets in blood vessels and recombined cells in bronchovascular cuff spaces but only very rare recombined cells and possible platelets in lymph. (**B**) *Prox1*-eGFP mice were given an o.a. dose of PKH26-PCL dye to label alveolar macrophages, then 5 days later mice were given o.a. LPS. Intravital imaging at 24 hours after LPS treatment showed labeling of alveolar macrophages (alv macs, asterisks) in alveoli but not in lymphatic vessels.

**Figure S3:**
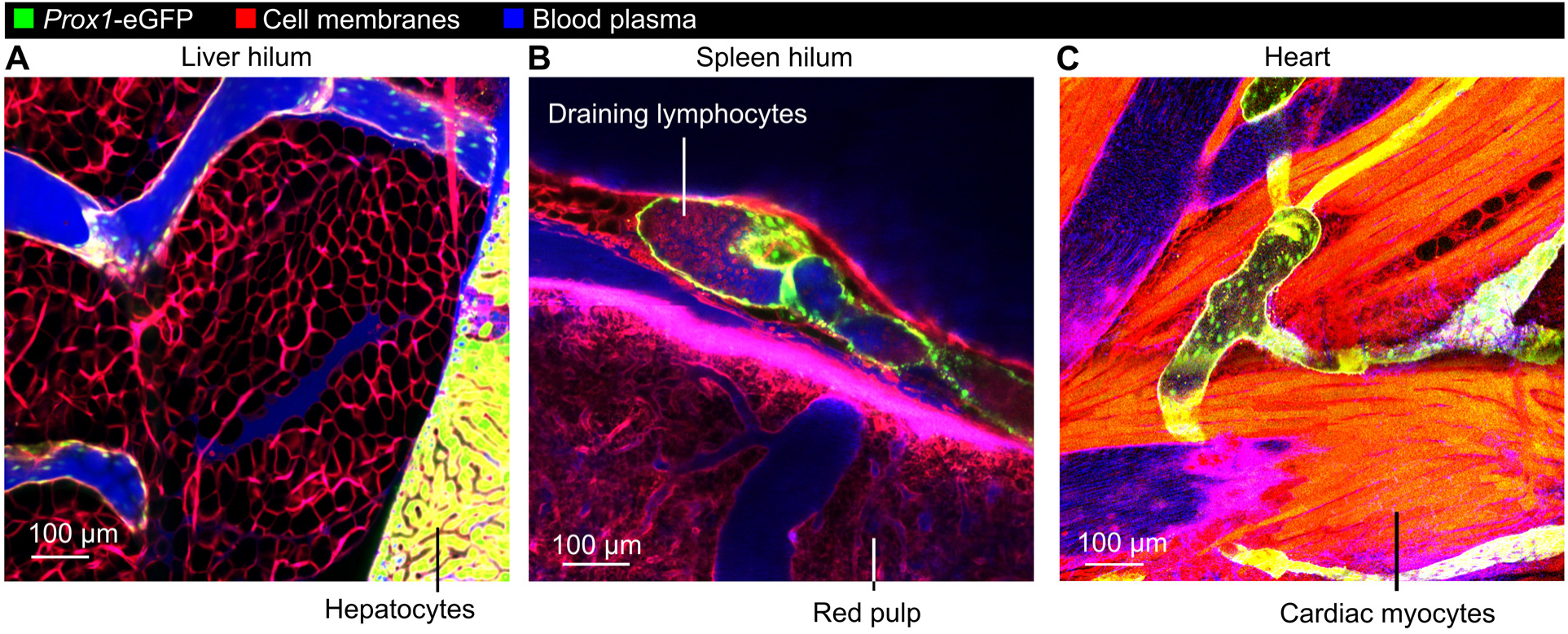
Stabilized imaging of lymphatic vessels draining the liver, spleen and heart. *Prox1*-eGFP:*Rosa26*^mTmG^ mice were given Evans blue i.v. prior to stabilized intravital imaging of: (**A**) the hilum of the liver; (**B**) the hilum of the spleen; and (**C**) the ventricular wall of the heart. Note free movement of Evans blue-labeled plasma proteins into liver and spleen draining lymphatics, likely due to the fenestrated endothelium lining blood vessels in these organs, as well as many leukocytes with lymphocyte morphology draining from the spleen, a secondary lymphoid organ.

